# Phylogenomic analyses using genomes and transcriptomes do not “resolve” relationships among major clades in Phrymaceae

**DOI:** 10.1101/2021.11.17.468996

**Authors:** Diego F. Morales-Briones, Nan Lin, Eileen Y. Huang, Dena L. Grossenbacher, James M. Sobel, Caroline D. Gilmore, David C. Tank, Ya Yang

## Abstract

**Premise of the study:** Phylogenomic datasets using genomes and transcriptomes provide rich opportunities beyond resolving bifurcating phylogenetic relationships. Monkeyflower (Phrymaceae) is a model system for evolutionary ecology. However, it lacks a well-supported phylogeny for a stable taxonomy and for macroevolutionary comparisons.

**Methods:** We sampled 24 genomes and transcriptomes in Phrymaceae and closely related families, including eight newly sequenced transcriptomes. We reconstructed the phylogeny using IQ-TREE and ASTRAL, evaluated gene tree discordance using PhyParts, Quartet Sampling, and cloudogram, and carried out phylogenetic network analyses using PhyloNet and HyDe. We searched for whole genome duplication (WGD) events using chromosome numbers, synonymous distance, and gene duplication events.

**Key results:** Most gene trees support the monophyly of Phrymaceae and each of its tribes. Most gene trees also support the tribe Mimuleae being sister to Phrymeae + Diplaceae + Leucocarpeae, with extensive gene tree discordance among the latter three. Despite the discordance, polyphyly of *Mimulus* s.l. is strongly supported, and no particular reticulation event among the Phrymaceae tribes is well supported. Reticulation likely occurred among *Erythranthe bicolor* and close relatives. No ancient WGD event was detected in Phrymaceae. Instead, small-scale duplications are among potential drivers of macroevolutionary diversification of Phrymaceae.

**Conclusions:** We show that analysis of reticulate evolution is sensitive to taxon sampling and methods used. We also demonstrate that genome-scale data do not always fully “resolve” phylogenetic relationships. They present rich opportunities to investigate reticulate evolution, and gene and genome evolution involved in lineage diversification and adaptation.

## INTRODUCTION

Phylogenomic analyses using genomes and transcriptomes often focus on “resolving” relationships that previously show conflict among markers or low statistical support using Sanger sequencing. With thousands of genes, these datasets are rich in insights beyond providing a bifurcating phylogenetic relationship, pointing to reticulate evolution, gene/genome duplications, and adaptation. However, detecting these events are difficult due to computational limitations, and studies often do not fully interrogate the data.

Monkeyflowers, as part of Phrymaceae, are a model system for evolutionary ecology (Wu et al., 2008; Twyford et al., 2015). With ca. 200 species, a primarily North American distribution, a rich history of ecological studies, and accumulating genomic resources, monkeyflower research has provided insights in speciation (Schemske and Bradshaw, 1999; Sobel, 2014), local adaptation (MacNair, 1983; Hall et al., 2010), pigment evolution (Streisfeld et al., 2013; Yuan et al., 2016; Ding et al., 2020), and development (Yuan, 2019). However, previous studies often focused on a single species or clades of closely related species (Stankowski and Streisfeld, 2015; Chase et al., 2017; Stankowski et al., 2019; Nelson et al., 2021), and we still lack a robust phylogenetic framework for the family. In addition, with phylogenetic uncertainty across Phrymaceae from previous analyses, the circumscription of *Mimulus* L. is under debate (Lowry et al., 2019; Nesom et al., 2019). Previous phylogenetic studies have established the polyphyly of *Mimulus* in its broad sense (Barker et al., 2012), and therefore we follow the narrow definition of *Mimulus* sensu Barker et al. (2012) that includes only seven species as part of the tribe Mimuleae, with the remaining species distributed into the genera *Diplacus* (tribe Diplaceae) and *Erythranthe* (tribe Leucocarpeae).

Previous molecular phylogenetic studies using Sanger sequencing have consistently supported five well-supported tribes within Phrymaceae [Fig. 1; the tropical Asian tribe Cyrtandromoeeae was only included in the sampling by Liu et al. (2020)]. However, phylogenetic relationships in Phrymaceae have been problematic due to discordance among analyses (Fig. 1) that used 1) different taxon sampling; 2) nuclear vs. plastome (cpDNA) markers, or even among cpDNA regions [*trnL-F* only (Beardsley and Olmstead, 2002) vs. six cpDNA regions (Liu et al., 2020)]; and 3) different analytical approaches (maximum likelihood vs. Bayesian, ITS + ETS, Liu et al., 2020). In addition, despite previous and ongoing whole genome and exome sequencing efforts (Hellsten et al., 2013; Edger et al., 2017; Nelson et al., 2021), there yet lacks multi-locus analyses using nuclear genes across major clades of the family.

**Figure 1.**
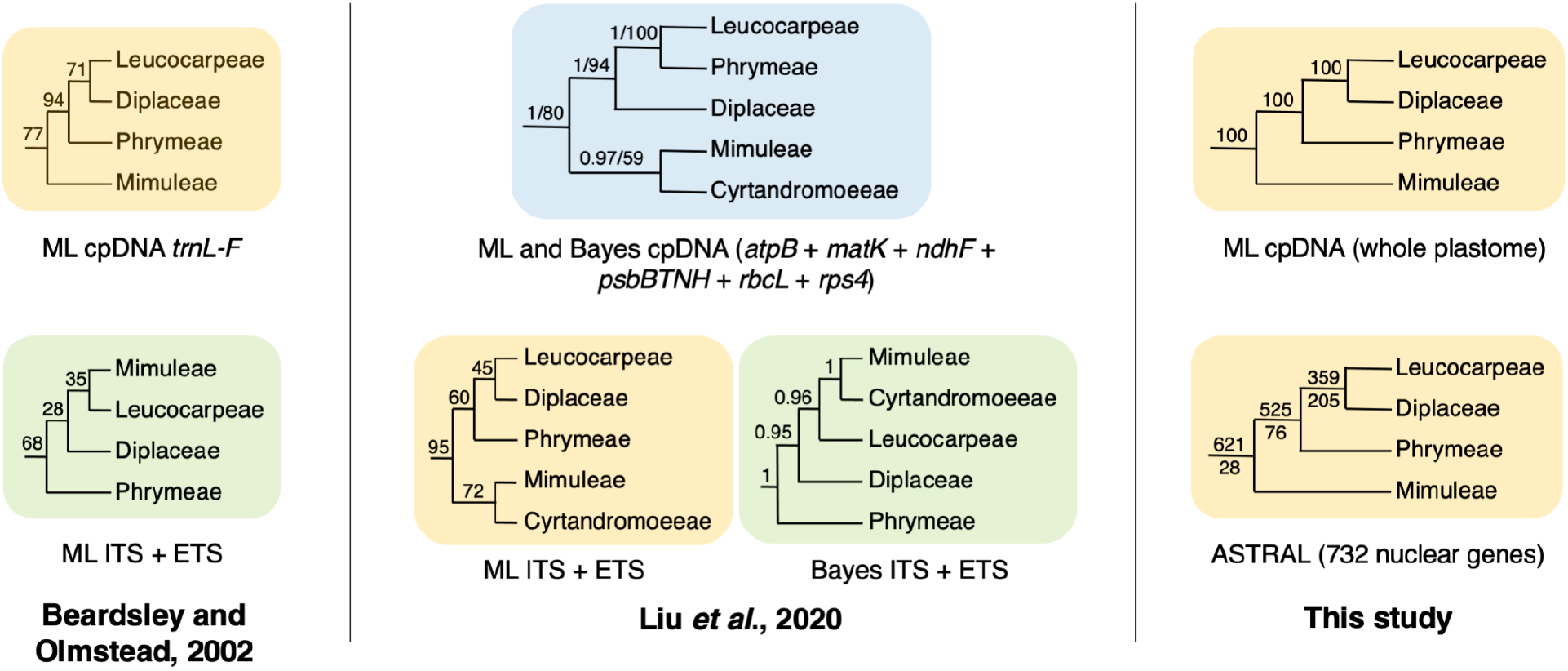
Summary of the topologies among the five tribes of Phrymaceae among studies focusing on the backbone relationship of the family. Numbers above branches are maximum likelihood (ML) bootstrap support (BS), Bayesian posterior probabilities (PP), or BS/PP. For the ASTRAL analysis, numbers above and below branches are the number of genes with concordant vs. discordant topologies that had MLBS >50 in each individual gene tree. Trees shaded by the same color share compatible backbone topologies.

Macroevolutionary analyses using a large number of nuclear genes are powerful not only for inferring the phylogenetic relationships and history of reticulate evolution, but also investigating gene and genome evolution associated with lineage diversification and adaptation. Previous investigation of patterns of chromosome number changes across the North American members of Phrymaceae suggested extensive polyploidy events across the family (Beardsley et al., 2004). However, comparison of linkage maps established that the higher chromosome base number in *Erythranthe lewisii* compared to *E. guttata* is due to chromosome fission and fusion instead of whole genome duplication (WGD; Fishman et al., 2014). To what extent WGD events are distributed, or the lack of, along the backbone of the family is still unexplored. Sampling of genomes and transcriptomes across major clades of Phrymaceae and analyzing both nuclear and cpDNA regions are needed to investigate the backbone structure in Phrymaceae and the genomic basis of the macroevolutionary diversification of Phrymaceae.

In this study we sampled transcriptomes and genomes covering four of the five tribes in Phrymaceae, including eight newly generated transcriptomes, to 1) provide a phylogenetic backbone and examine gene tree discordance and 2) investigate patterns of gene and genome duplication in the family. We found that most gene trees support Mimuleae being sister to Phrymeae+Diplaceae+Leucocarpeae, with the relationship among the latter three showing extensive gene tree discordance. However, no particular reticulation event among the Phrymaceae tribes was strongly supported. Instead, we found evidence for reticulation events among closely related species contributing to *Erythranthe bicolor*. Our analyses did not support any ancient WGD events in Phrymaceae; instead, small-scale gene duplications in defense, stress response, growth and development, and certain biochemistry pathways are candidates for potential drivers underlie macroevolutionary diversification of Phrymaceae.

## MATERIALS AND METHODS

### Taxon sampling

Transcriptomes were newly generated from eight accessions representing seven ingroup Phrymaceae species for this study (Appendix S1). Seeds were collected from natural populations and were cold-treated in soil for a week at 4°C in the dark before growing with 15-hour daylight in a greenhouse. Young leaves and flower buds were flash frozen in liquid nitrogen. RNA extraction, library preparation (rRNA removal or poly-A enrichment), and sequencing procedures are detailed in Appendix S1. In addition, we included three genomes and five transcriptomes from Phrymaceae that are publicly available (Appendix S2). Together our taxon sampling included 16 datasets representing 15 Phrymaceae species in four of the five tribes (missing the tropical Asian tribe Cyrtandromoeeae) and five of the 14 genera (Barker et al., 2012; Liu et al., 2020). We also included four genomes and four transcriptomes in closely related Lamiales families (Zhang et al., 2020).

### Data processing for nuclear genes

Read processing, assembly, and translation were carried out following Morales-Briones et al. (2021). Homology inference started with an all-by-all BLASTN search on coding sequences (CDS) with an *E* value cutoff of 10. Hits were filtered with a minimal hit coverage of 40%. Homolog groups were clustered using MCL v. 14-137 (van Dongen, 2000) with a minimum minus log-transformed *E* value cutoff of 5 and an inflation value of 1.4. Only clusters with at least 20 out of 24 taxa were retained. Sequences from each cluster were aligned using the OMM_MACSE pipeline v. 10.02 (Scornavacca et al., 2019), which pre-filters non-homologous sequence fragments with HMMCleaner (Di Franco et al., 2019) before translation accounting for frameshifts using MACSE v. 2.03 (Ranwez et al., 2018). The resulting CDS alignments were trimmed to remove columns with more than 90% missing data using Phyx (Brown et al., 2017). Homolog trees were built with RAxML v. 8.2.11 (Stamatakis, 2014) using the GTRCAT model and 200 rapid bootstrap (BS) replicates. Monophyletic and paraphyletic tips of the same dataset were removed, keeping the tip with the highest number of characters in the trimmed alignment. Spurious tips were detected and removed with TreeShrink v. 1.3.2 (Mai and Mirarab, 2018) with the ‘per-gene’ mode and a false positive error rate threshold (α) of 0.001. The resulting trees were visually inspected, and deep paralogs producing internal branch lengths longer than 0.25 were cut apart retaining subclades with a minimum of 20 taxa to obtain final homolog trees.

Orthology inference was carried out using the ‘monophyletic outgroup’ approach from Yang and Smith (2014). The approach filters unrooted homolog trees, requiring outgroups to be monophyletic and single-copy. It then traverses the ingroup clade rooted by outgroups from root to tip and trims the side with less taxa each time a gene duplication event was detected, until every taxon is represented by a single sequence. We set the three Lamiaceae genomes as outgroups (Zhang et al., 2020), keeping ortholog groups that included at least 15 taxa.

### Species tree inference and evaluation of support

Sequences from individual ortholog groups were aligned using OMM_MACSE. Columns with more than 20% missing data were trimmed with Phyx, and only alignments with at least 1000 characters and all 24 taxa were retained. We first estimated a maximum likelihood (ML) tree of the concatenated matrix with IQ-TREE v. 2.1.13 (Minh et al., 2020) searching for the best partition scheme (Lanfear et al., 2012), followed by ML tree inference and 1000 ultrafast bootstrap replicates. To estimate a coalescent-based species tree, we first inferred individual gene trees with IQ-TREE using extended model selection (Kalyaanamoorthy et al., 2017) and 200 non-parametric bootstrap replicates for clade support. Gene trees were then used to infer a species tree with ASTRAL-III v. 5.6.3 (Zhang et al., 2018) using local posterior probabilities (LPP; Sayyari and Mirarab, 2016) to assess clade support.

To explore discordance among gene trees, we calculated the number of concordant and discordant bipartitions on each node of the species tree using PhyParts (Smith et al., 2015). We mapped bipartitions from gene trees with local BS support of at least 50% against the IQ-TREE tree from the concatenated supermatrix (identical to the ASTRAL tree; see results). Next, to distinguish conflict from poorly supported branches, we carried out a Quartet Sampling (QS; Pease et al., 2018) analysis using the concatenated supermatrix, the IQ-TREE tree, and 1000 replicates. Lastly, to visualize gene tree conflict, we built a cloudogram using the DensiTree function of phangorn v. 2.7.1 (Schliep et al., 2017). Individual ortholog gene trees were time-calibrated for visualization purposes with TreePL v. 1.0 (Smith and O’Meara, 2012). The root was fixed to 70.3 MYA, the most recent common ancestor (MRCA) of Lamiaceae was fixed to 57.69 MYA, and the MRCA of all remaining species was fixed to 68.8 MYA based on Zhang et al. (2020).

### Plastome assembly and tree inference

We obtained nine reference plastomes from RefSeq (Appendix S3). For the remaining species, we carried out plastome assemblies from either transcriptomic or genomic reads (Appendix S3) with Fast-Plast v. 1.2.8 (McKain and Wilson, n.d.). In four cases, those assemblies resulted in low plastome coverage and were redone using alternative transcriptomic or genomic libraries (Appendix S3). When the resulting plastome were incomplete (7 out of 14 accessions), filtered contigs from Spades v. 3.9.0 (Bankevich et al., 2012) were mapped to the closest available reference plastome using Geneious v. 11.1.5 (Kearse et al., 2012) to produce oriented and contiguous contigs with missing regions masked with ‘N’. The assembly of *Striga asiatica* had many contigs that were poorly mapped even to congeneric plastomes, likely due to major structural rearrangements in this hemiparasitic species of Orobanchaceae (Frailey et al., 2018). We replaced it with the published plastome of *Striga forbesii* (Appendix S3) for downstream analyses.

The resulting plastomes with one Inverted Repeat removed were aligned with MAFFT and columns with more than 50% missing data were trimmed with Phyx. An ML tree was inferred with IQ-TREE with automated extended model selection and 1000 rapid BS replicates. Additionally, we used QS with 1,000 replicates to evaluate branch support.

### Testing potential reticulation at nodes with elevated levels of gene tree conflict

We investigated two areas with elevated gene tree conflict: 1) the backbone of Phrymaceae using one species for each well-supported clade corresponding to a tribe; and 2) among *Erythranthe cardinalis, E. lewisii*, and *E. bicolor*.

For each of the two areas, we first ran PhyParts using all taxa from the reduced dataset. We then removed one taxon at a time to determine which taxon produced the highest gene tree conflict. We inferred species networks using ML (Yu et al., 2014) in PhyloNet v. 3.6.9. (Than et al., 2008) with the command “InferNetworks_ML’’ from individual ML gene trees. Network searches were performed allowing for up to three reticulation events, and optimizing the branch lengths and inheritance probabilities of the inferred species networks. To estimate the optimal number of reticulations and to test whether a species network fits our gene trees better than a strictly bifurcating tree, we computed the likelihood scores of the nuclear and plastid trees given the individual gene trees using the command ‘CalGTProb’ (Yu et al., 2012). We performed model selection using the Akaike information criterion (AIC; Akaike, 1973), bias-corrected AIC (AICc; Sugiura, 1978) and the Bayesian information criterion (BIC; Schwarz, 1978). Next, we performed a more thorough Phylonet analysis using a Bayesian inference of species networks approach (Wen et al., 2016) with the command “MCMC_GT”, full likelihood, and allowing up to three recticulations events. Analyses consisted of four independent runs with four reversible-jump Markov chain Monte Carlo (RJMCMC) chains, temperatures set to one cold and two hot chains (1.0, 2.0 and 3.0 respectively), 30 million generations, sampling every 1000 generation, and a burn-in of 500,000 generations. The four MCMC runs were summarized with the command “MCMC_GT -sum”. Convergence was assessed once the posterior sampling reached ESS (effective sample size) ≥ 200.

In addition to PhyloNet, we also tested for hybridization with HyDe (Blischak et al., 2018) that uses site pattern frequencies (Kubatko and Chifman, 2019) to quantify admixture (γ) between two parental lineages that form a hybrid lineage. We tested all triplet combinations in all directions using ‘run_hyde.py’, the concatenated nuclear alignment, and a mapping file to assign individuals to species. Test significance was assessed with a Bonferroni correction (α= 0.05) for the number of tests conducted with estimates of γ between 0 and 1 (Blischak et al., 2018).

### Gene and whole genome duplication events

We employed three approaches (Yang et al., 2018) to detect WGD events in Phrymaceae. 1) We summarized chromosome counts from Nesom (2012) and the Chromosome Counts Database (Rice et al., 2015). 2) We mapped gene duplication events onto the inferred species tree by extracting rooted ingroup clades from the final homolog trees by requiring an average BS ≥ 50 and at least 15 taxa. Gene duplication events were then mapped onto the MRCA on the species tree when two or more taxa overlapped between the two daughter clades on the rooted ingroup clade (“extract_clades.py” and “map_dups_mrca.py” from https://bitbucket.org/blackrim/clustering). 3) We analyzed the distribution of synonymous distances (Ks) from RNA-seq (https://bitbucket.org/blackrim/clustering; “ks_plots.py”). Ks peaks were identified using a mixture model in mixtools v. 1.2.0 (Benaglia et al., 2009).

To explore genes with elevated instances of gene duplication within Phrymaceae, we extracted Phrymaceae clades from the final homologs. We then obtained functional annotation for the top 10 Phrymaceae clades with the highest number of genes using the *Erythranthe guttata* genome annotation (Hellsten et al., 2013).

## RESULTS

### Sequence processing

Organellar reads represented 30–57% of quality-filtered read pairs in RNA-seq libraries prepared using RiboZero, compared to 0.13–0.3% in libraries prepared using Poly-A enrichment (Appendix S1). Of the eight newly generated transcriptomes, we retained 13.7–21.4 million nuclear read pairs after quality filtering and separating organellar reads; each CDS set represented 49–62% of nuclear genes when compared against the *Erythranthe guttata* reference genome (Appendix S2). Although libraries prepared with RiboZero produced lower numbers of nuclear reads, they produced more contiguous assemblies and some of the highest numbers of genes in our final nuclear ortholog set (Appendix S2). This can be due to ribosomal depletion resulting in more even coverage in slightly degraded transcripts compared to poly-A enrichment.

Assemblies from each of the six genomic libraries produced full plastomes in a single contig (Appendix S3). Despite the large numbers of plastid reads from the six RiboZero RNA libraries, only one complete plastome was assembled due to uneven coverage. Still, RiboZero RNA libraries produced contiguous contigs that covered most of the plastomes and recovered similar numbers of CDS compared to the full plastomes (Appendix S3).

### Orthology inference and phylogenetic analysis

The final set of nuclear orthologs included 732 genes, and the concatenated matrix consisted of 1,246,075 aligned columns with a character occupancy of 93.1% (Appendix S2). The topologies from the IQ-TREE and ASTRAL trees were identical and all nodes had maximum support (BS =100, LPP = 100; Figs. 1, 2A, & Appendix S4). The monophyly of Phrymaceae and each tribe of Phrymaceae was strongly supported by most informative gene trees and full QS support (1/–/1; i.e., all sampled quartets supported that branch; blue in Fig. 2A and Appendix S5). The sister relationship of Leucocarpeae and Diplaceae was supported by 359/564 informative gene trees and strong Quartet Concordance (QC = 0.3), but the Quartet Differential (QD = 0) indicates the presence of a single alternative topology (Diplaceae sister to Phrymeae). Similarly, the placement of Phrymeae was supported by 525/601 informative gene trees, strong QC (0.72) and signal of a single potential alternative topology (QD = 0; Mimulaeae sister to Phrymeae). The cloudogram (Fig. 2C) showed discordance in the backbone of Phrymaceae, especially on the placement of Phrymeae (“Phrlep”), consistent with the concordance analyses and QS results.

**Figure 2.**
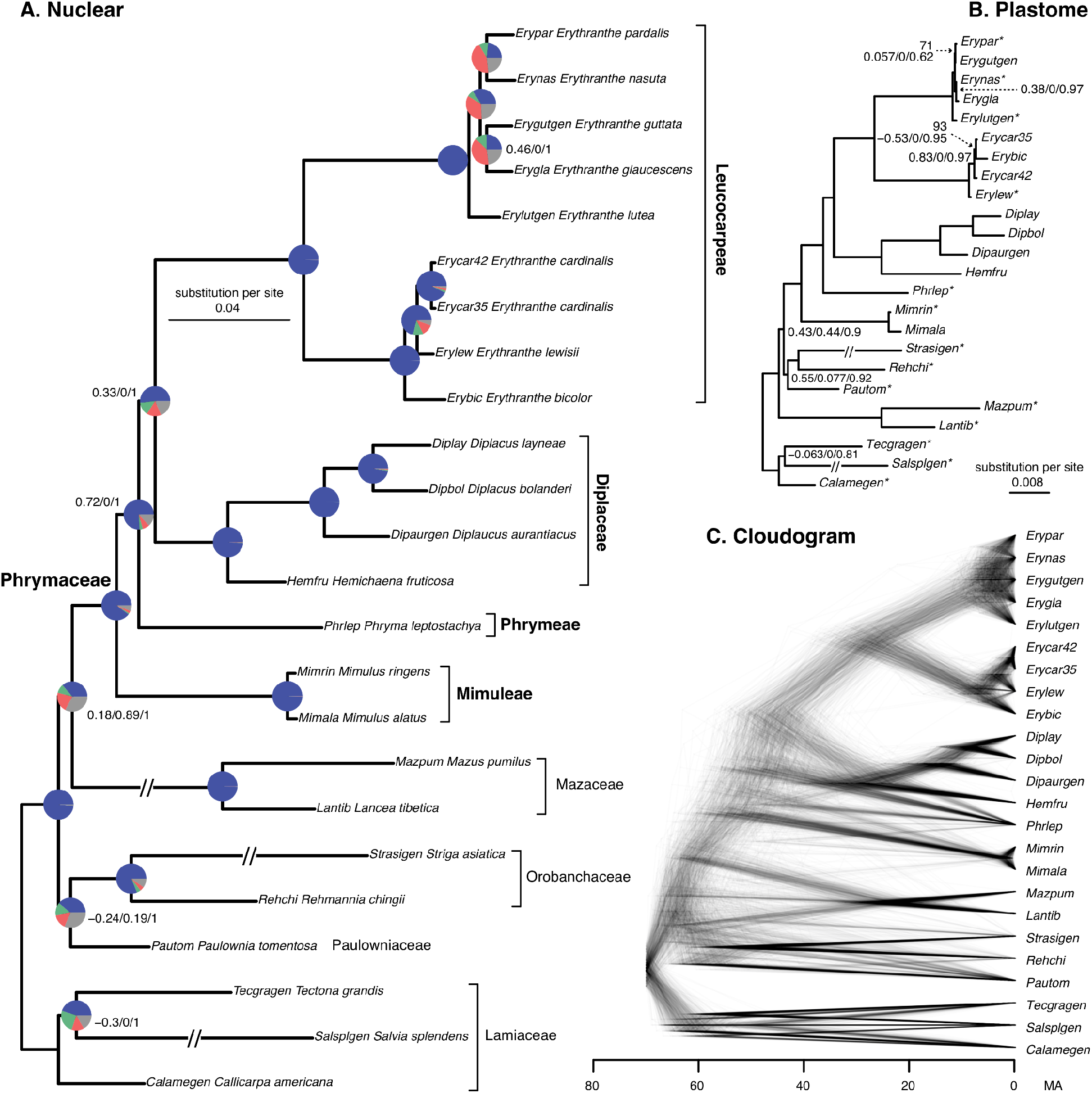
Maximum likelihood phylogeny of Phrymaceae inferred with IQ-TREE from the concatenated 732-nuclear gene supermatrix. Quartet Sampling (QS) scores are shown below branches. QS scores: Quartet concordance/Quartet differential/Quartet informativeness. Nodes with maximum QS support (1/—/1) not shown. All nodes have full bootstrap support (BS =100) and local posterior probability (LLP =1). See Appendix S4. Pie charts represent the proportion of ortholog trees that support that clade (blue), the main alternative bifurcation (green), the remaining alternatives (red), and the remaining with (conflict or support) <50% bootstrap support (gray). Branch lengths are in number of substitutions per site (scale bar). Exceptionally long branches are shortened with a broken segment (//) for illustration purposes (See Appendix S5 for original branch lengths); b) Maximum likelihood phylogeny inferred with IQ-TREE from plastomes. Bootstrap support shown above branches and QS scores below the branches. Full BS and QS support values are not shown. Branch lengths are in number of substitutions per site (scale bar). Longest branches are shortened with a broken segment (//) for illustration purposes (See Appendix S5 for original branch lengths); c) Cloudogram inferred from 732 nuclear ortholog trees. Scale in millions of years ago (MA).

The final cpDNA alignment included 128,056 characters with a character occupancy of 89%. The plastome topology recovered the monophyly of Phrymaceae and each of its tribes with maximum support (BS =100, QS 1/-/1; Fig. 2B) and the backbone relationships were identical to the nuclear results (Fig. 2A, Appendix S5). However, relationships among closely related ingroup taxa differ in two places (Fig. 2, A vs. B): 1) The two *Erythranthe cardinalis* accessions were sister to each other in the nuclear tree but were paraphyletic with *E. bicolor* nested among them in the cpDNA tree; 2) relationships among *Erythranthe pardalis, E. nasuta, E. guttata*, and *E. glaucescens* showed extensive gene tree conflict among nuclear genes, low quartet concordance, and dominant secondary topologies in the cpDNA tree, and conflicting topology between cpDNA and nuclear trees. In addition, extensive nuclear gene tree conflict and discordance between nuclear and cpDNA trees are present among other Lamiales families sampled.

### Phylogenetic network analyses

Phylogenetic network analyses focused on two ingroup areas with elevated levels of conflict. To investigate the backbone of Phrymaceae, we used one taxon to represent each tribe. The cloudogram (Fig. 2C) showed *Phryma leptostachya* (Phrymeae) shifted its placement among other Phrymaceae tribes. When removing one tip at a time, conflict among nuclear gene trees (red and green in Fig. 3A) reduced the most when removing *Phryma*, followed by removing *Erythranthe guttata* (Leucocarpeae). Visual inspection of individual gene trees and alignments confirmed that *Phryma*’s placement shifted among gene trees with short internal branches attached to the backbone of Phrymaceae. PhyloNet ML searches (Appendix S6) recovered three networks with small amounts of gene flow towards *Mimulus ringens*. Model selection (Appendix S7) using AIC and AICc both preferred three reticulations while BIC did not support significant differences among the three networks. MPP from the MCMC PhyloNet searches (Fig. 3A) recovered the same network as the 1-reticulation network from ML, and estimated that 9.17% of *M. ringens* genes had contribution from *E. guttata*, with the 95% credible set consisted of a single network. HyDe (Appendix S8) results indicated *E. guttata* and *Diplacus aurantiacus* were each derived from reticulation between the stem lineage of Mimuleae and Phrymeae, and *E. guttata* from reticulation between Mimuleae and Diplaceae. As almost all informative gene trees supported the monophyly of each Phrymaceae tribe, any potential hybridization events would have occurred among stem branches of tribes. Given the disagreement among PhyloNet and HyDe analyses, none of the putative hybridization events were well supported. Phylogenetic uncertainty, ILS, and population structure likely contributed to the high levels of gene tree discordance among Phrymaceae tribes.

**Figure 3.**
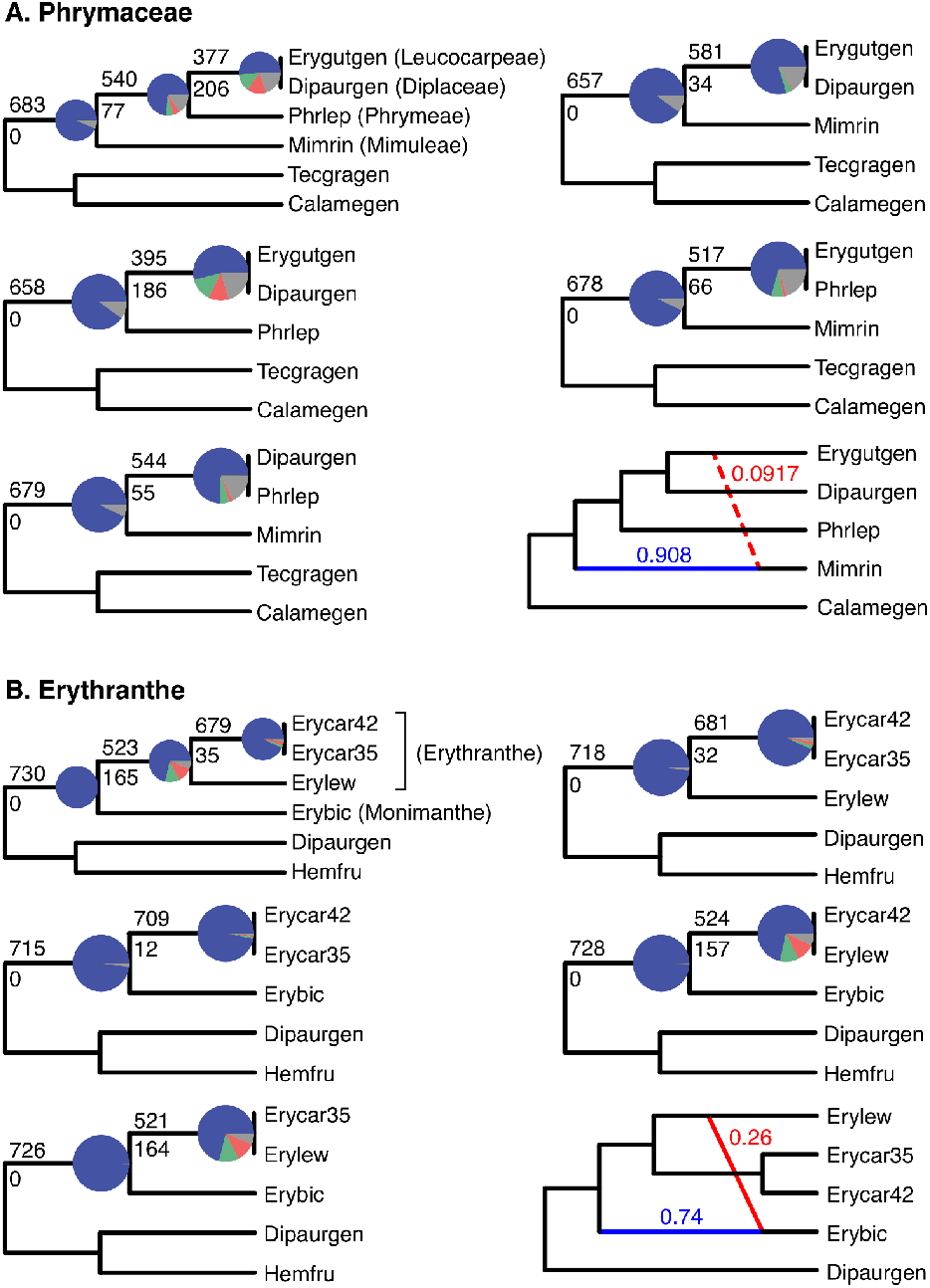
Conflict and network analyses using reduced datasets. Cladograms showing relationships among the reduced dataset, removing one tip at a time, and Maximum posterior probability (MPP) network from the PhyloNet Bayesian inference. A) Phrymaceae backbone with one representative species for each tribe. B) *Erythranthe cardinalis, E. lewisii*, and *E. bicolor*. Pie charts on cladograms represent the proportion of gene trees that support that clade (blue), the main alternative bipartition (green), the remaining alternatives (red), and conflict or support with <50% bootstrap support (gray). Number above and below branches represent the number of concordant and discordant gene trees, respectively. Red and blue branches in networks indicate the minor and major edges respectively, of hybrid nodes, with the inheritance probabilities next to each branch. Dotted edge indicates uncertainty in inference (see Results).

The second instance of conflict we focused on was among *Erythranthe bicolor, E. cardinalis*, and *E. lewisii*. Consistent with the cloudogram (Fig. 2C) where some gene trees supported *E. bicolor* being sister to *E. lewisii*, removing either *E. bicolor* or *E. lewisii* (Fig. 3B) removed most of the gene tree conflicts. PhyloNet ML analysis (Appendix S6) recovered networks with 38–48% of *E. bicolor* genes from *E. lewisii* or its close relatives. AIC or AICc did not prefer any network, while BIC preferred the 1-reticulation network (Appendix S7). PhyloNet MCMC searches recovered a 95% credible set of three 1-reticulation networks with gene flow towards *E. bicolor*, but the source of gene flow varied among networks (Appendix S9). The MPP (54% of the credible set) showed that *E. bicolor* had 26% genes from *E. lewisii*, similar to the 1-reticulation ML network (Fig. 3B). HyDe, on the other hand, did not recover any significant hybridization events.

Although we also detected extensive gene tree conflict among *Erythranthe pardalis, E. nasuta, E. guttata*, and *E. glaucescens*, as branches among them were short and we lacked any intraspecific sampling, we did not carry out additional analyses on reticulate evolution.

### Gene and Genome Duplications

Mapping gene duplication events did not reveal any node with more than 4.3% of gene duplications in Phrymaceae (Appendix S10). Similarly, the Ks plots (Appendix S11) did not support any Phrymaceae-specific WGD that involved more than one species sampled. All 24 datasets included in this study shared two optimal mixing components (i.e. Ks peaks). The first component had a Ks mean of 1.8–2.2, corresponding to a whole-genome triplication event early in the core eudicots (Jiao et al., 2012). The second component had Ks means of 0.3–0.9 (lower in woody species and higher in herbaceous species), corresponding to a WGD at the MRCA of the core Lamiales (Zhang et al., 2020). A third component at Ks∼0.1 was found only in *Erythranthe lutea*, corresponding to a previously reported WGD event (Edger et al., 2018). In addition, *Diplacus layneae* and *Erythranthe guttata* each showed a putative Ks peak at ∼0.04 and 0.09 respectively. However, chromosome counts (Appendix S10) did not support a WGD in either species. Overall, all sampled Phrymaceae species except *Erythranthe lutea* had low chromosome counts compared to outgroups, and the uptick in Ks density below 0.1 was likely due to recent small-scale duplications. Among non-Phrymaceae species, *Salvia splendens* showed a Ks peak ∼0.1, consistent with a WGD in *Salvia* and relatives within Lamiaceae (Godden et al., 2019). Both *Mazus pumilus* and *Lancea tibetica* showed a Ks peak ∼0.07, which could be due to WGD or small-scale duplications.

The ten Phrymaceae genes (Appendix S12) with the highest numbers of copies were involved in defense/immune response (Serine protease inhibitor, aspartyl protease, MLP-like protein), stress response (HSP20-like chaperones, Ribosomal protein L10 family protein), mitochondria organization (prohibitin 2), regulating plant growth (small auxin up-regulated RNA-like auxin-responsive protein family), cell wall architecture (Glycosyl hydrolase), and various other biochemical processes (S-adenosyl-L-methionine-dependent methyltransferases, hydroxymethyltransferase 4). Since we reduced sequences from the same sample that formed monophyletic or paraphyletic groups, we ignored isoforms from alternative splicing, assembly artifacts, and recent copy number increase involving only a single sample, as these are difficult to quantify using *de novo* assembled transcriptomes. Therefore, only gene duplication events involving more than two taxa in our sampling contributed to our copy number counts. Given our much denser taxon sampling in Leucocarpeae and Diplaceae, the top ten are heavily influenced by genes that had multiple rounds of gene duplications in these two tribes.

## DISCUSSION

### Extensive gene tree discordance and potential hybridization events in Phrymaceae

Our phylogenomic analyses recovered strong support of the monophyly of Phrymaceae and each of its tribes sampled. We also recovered extensive and well-supported gene tree discordance along the backbone of Phrymaceae. The discordance is not an artifact of gene and genome duplications; nor is any particular reticulation event well supported by phylogenetic network analyses and hypothesis testing. Therefore, phylogenetic uncertainty, ILS, and population structure likely contributed to the extensive gene tree discordance among Phrymaceae tribes.

Among closely related species, our phylogenetic network analyses support introgression from *E. lewisii* or close relatives towards *E. bicolor*. In addition, our plastome analysis (Fig. 2B) recovered *E. bicolor* being nested among accessions of *E. cardinalis*, suggesting that *E. cardinalis* is also involved in the reticulation. However, without sampling of other closely related species or additional within-species sampling, the timing, source, and prevalence of the introgression is unclear. Nelson et al. (2021) analyzed over 8,000 nuclear gene trees and recovered extensive reticulation among *E. lewisii, E. cardinalis*, and *E. parishii* (not sampled in our study), with *E. bicolor* set as the outgroup. Our analyses suggest that additional clades are involved in reticulations involving *E. lewisii, E. cardinalis*, and/or other close relatives.

In summary, our phylogenetic analyses suggest that: 1) network inferences are sensitive to the methods used, sources of data, and taxon sampling, including the choice of both ingroups and outgroups. 2) Despite phylogenetic uncertainty along the Phrymaceae backbone, most gene trees support Mimuleae (likely together with the unsampled Cyrtandromoeeae) being sister to a strongly supported clade of Phrymeae + Diplaceae + Leucocarpeae, consistent with previous Sanger-based studies (summarized in Barker et al. 2012). 3) Our results strongly support the polyphyly of the monkeyflower genus *Mimulus* s.l. (= part of Mimuleae + part of Leucocarpeae + part of Diplaceae).

### Genomic drivers of macroevolution in Phrymaceae

Analyses of chromosome counts, gene tree mapping, and Ks plots did not find any evidence for ancient WGD in Phrymaceae that involves more than one species in our taxon sampling. Our results are consistent with the previous comparisons using linkage maps between *E. lewisii* and *E. guttata* (Fishman et al., 2014) and whole genome sequences *E. guttata* and *E. lutea* (Edger et al., 2018), both primarily focused on species in the tribe Leucocarpeae. Our analysis broadened the genome-wide sampling to four of the five tribes in Phrymaceae and found that ancient WGD is not a driving force in macroevolution of Phrymaceae. Instead, reticulate evolution, small-scale gene duplication in genes involved in defense, stress response, growth and development, and additional biochemical pathways are some of the potential drivers of macroevolutionary diversification in Phrymaceae. Our study provides initial insights into the gene space of species across Phrymaceae and potential genomic drivers of macroevolution in the family. In addition, our newly generated transcriptome datasets using RiboMinus provide data for future studies looking into non-coding RNAs.

## CONCLUSIONS

Our phylogenomic analysis evaluated the support (or the lack of) in the backbone of Phrymaceae, confirmed the polyphyly of *Mimulus* s.l., and detected an area of reticulate evolution among closely related species. We show that analysis of reticulate evolution is sensitive to taxon sampling and methods used. We also show a lack of ancient WGD events in Phrymaceae; instead, small-scale gene duplications are potential drivers underlie macroevolutionary diversification of Phrymaceae.

Our analyses demonstrate that genome-scale data do not always “resolving” phylogenetic relationships. Instead, they provide resolution for some areas, but also recover “clouds” and “networks” that point to future opportunities for investigating their significance in adaptation and lineage diversification.

## Acknowledgments

The authors thank Rahul Roy for assistance with greenhouse work, and for Aaron Lee and Benjamin Cooper for commenting on the draft. Funding was provided by the University of Minnesota to DFM-B, YH, and YY; China Scholarship Council to NL (CSC: 201904910676), National Science Foundation grant to JMS (DEB-1856158), and the Frost Fund by the California Polytechnic State University to DLG.

## Author Contributions

YY and DLG designed the study; YH, DLG, JMS, CDG, and YY generated the data; DFM-B, NL, and YY analyzed data and drafted the manuscript. All authors edited the manuscript and approved the final version.

## Data Availability Statement

Raw reads of newly sequenced transcriptomes were deposited in the NCBI Sequence Reads Archive (BioProject: PRJNA770153). Analysis files are available from Dryad (to be deposited).

## Supporting Information

Additional supporting information may be found online in the Supporting Information section at the end of the article.

**Appendix S1**. Collection, plant growth, and sequencing information for the eight newly generated transcriptomes.

**Appendix S2**. Taxon sampling, source of data, and nuclear matrix statistics. Naming authorities above species level (Stevens, n.d.): 1) Order: Lamiales Bromhead. 2) Lamiales families: Phrymaceae Schauer, Orobanchaceae Ventenat, Mazaceae Reveal, Paulowniaceae Nakai, Orobanchaceae Ventenat. 3) Phrymaceae tribes: Diplaceae Bo Li, B. Liu, S. Liu & Y. H. Tan; Phrymeae Hogg; Leucocarpeae Conzatti; Mimuleae Dumortier; Cyrtandromoeeae Bo Li, B. Liu, S. Liu & Y. H. Tan. 4) Phrymaceae genera: *Diplacus* Nuttall, *Hemichaena* Bentham, *Erythranthe* Spach, *Mimulus* L., *Phryma* L.

**Appendix S3**. Plastome assembly and sources information.

**Appendix S4**. A. Maximum likelihood phylogeny of Phrymaceae inferred with IQ-TREE from the concatenated 732-nuclear gene supermatrix. Numbers above branches represent bootstrap support (BS). Branch lengths are in number of substitutions per site (scale bar on the bottom). B. ASTRAL-Pro tree of Phrymaceae inferred from the 732 nuclear gene trees. Local posterior probabilities (LLP) are shown next to nodes. Internal branch lengths are in coalescent units (scale bar on the bottom). C. Maximum likelihood phylogeny of Phrymaceae inferred with IQ-TREE from plastomes. BS values are shown above branches. Branch lengths are in number of substitutions per site (scale bar on the bottom)

**Appendix S5**. Maximum likelihood cladogram of Phrymaceae inferred with IQ-TREE from the concatenated 732-nuclear gene supermatrix. Pie charts represent the proportion of gene trees that support that clade (blue), the main alternative bifurcation (green), the remaining alternatives (red), and conflict or support that have <50% bootstrap support (gray). Number above and below branches represent the number of concordant and discordant informative gene trees, respectively.

**Appendix S6**. Species network inferred from PhyloNet maximum likelihood analyses with one to three maximum reticulations of the reduced data sets. A) Phrymaceae backbone. B) *Erythranthe cardinalis, E. lewisii*, and *E. bicolor*. Red and blue branches indicate the minor and major edges, respectively, of hybrid nodes. Numbers next to curved branches indicate inheritance probabilities for each hybrid node.

**Appendix S7**. Model testing between trees and PhyloNet networks for the reduced data sets of Phrymaceae and *Erythranthe*. The number of parameters were set to equal the number of branch lengths plus the number of inheritance probabilities. The number of gene trees was used to correct for finite sample size.

**Appendix S8**. HyDe tests for hybridization events along the backbone of Phrymaceae.

**Appendix S9**. Phylonet Bayesian inference 95% credibility set for the reduced data set of *Erythranthe cardinalis, E. lewisii*, and *E. bicolor*. Red and blue branches indicate the minor and major edges, respectively, of hybrid nodes. Numbers next to curved branches indicate inheritance probabilities for each hybrid node.

**Appendix S10**. Maximum likelihood cladogram of Phrymaceae inferred with IQ-TREE from the concatenated 732-nuclear gene supermatrix. Numbers above branches are gene duplication counts and numbers below branches are gene duplication percentages. Numbers next to species names are gametophytic chromosome counts. All chromosome counts are from the Chromosome Counts Database (Rice et al., 2015), except *Erythranthe pardalis* (Nesom, 2012). When multiple independent counts gave a single consistent chromosome number but different counts were each reported by a single study, we ignored the outlier numbers. Inset: Histogram of percentages of gene duplication per branch.

**Appendix S11**. Distribution of synonymous distance among gene pairs (Ks plots) for each genome or transcriptome. A) Plots of raw Ks distances between 0 and 3. B) Plots of Ks distances zooming in to values between 0 and 0.5. C) Plots of log-transformed Ks distances. Colored lines indicate components inferred using a mixture model. Blue lines indicate a component from an ancestral whole genome triplication event early in core eudicots; red lines are from more recent whole genome or small-scale duplication events.

**Appendix S12**. Phrymaceae clades extracted from the final homologs with the highest number of copies.

