## Supplementary figures and images for "Phylogenomic analyses using genomes and transcriptomes do not “resolve” relationships among major clades in Phrymaceae"

### Appendix S4

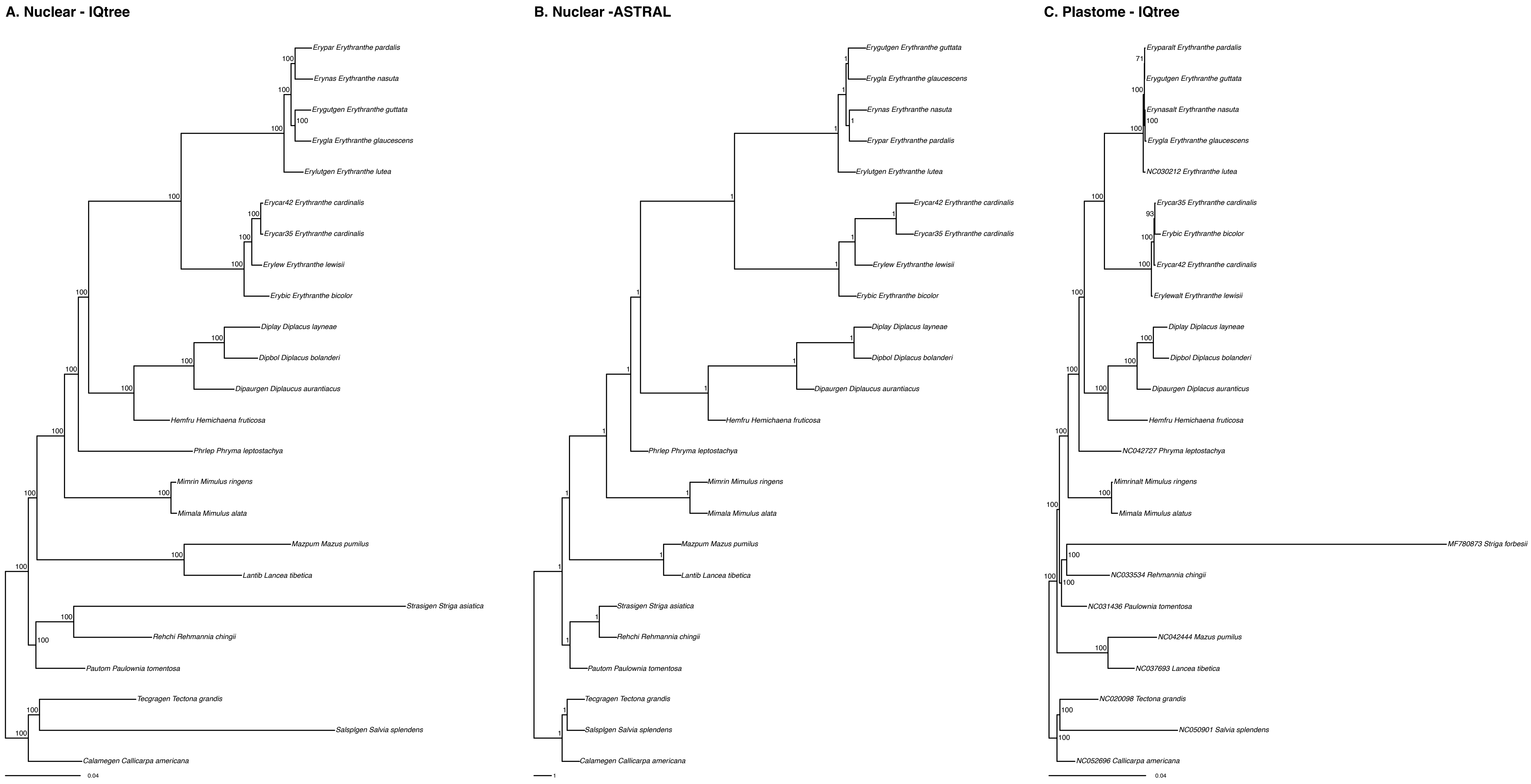

### Appendix S5

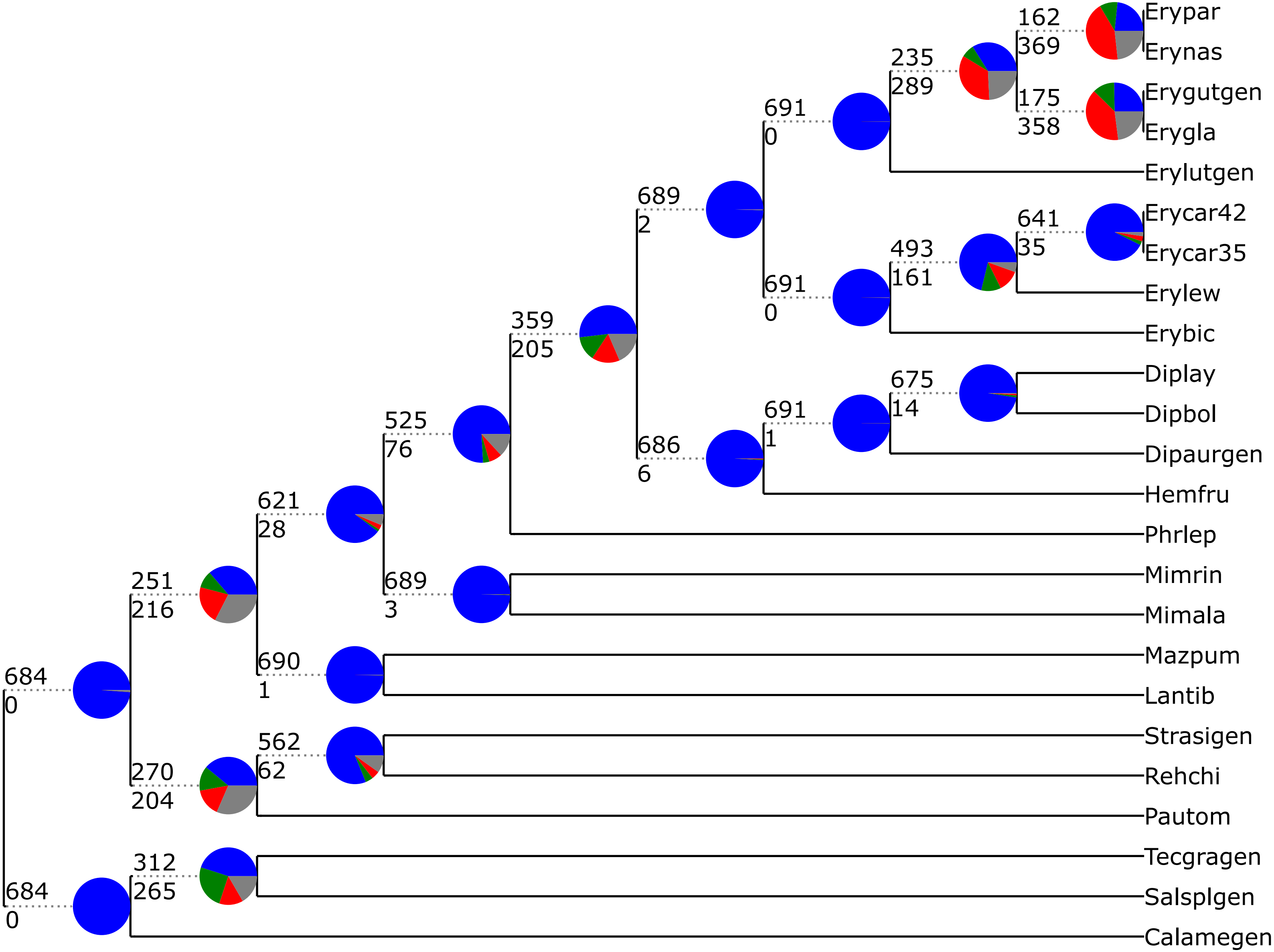

### Appendix S6

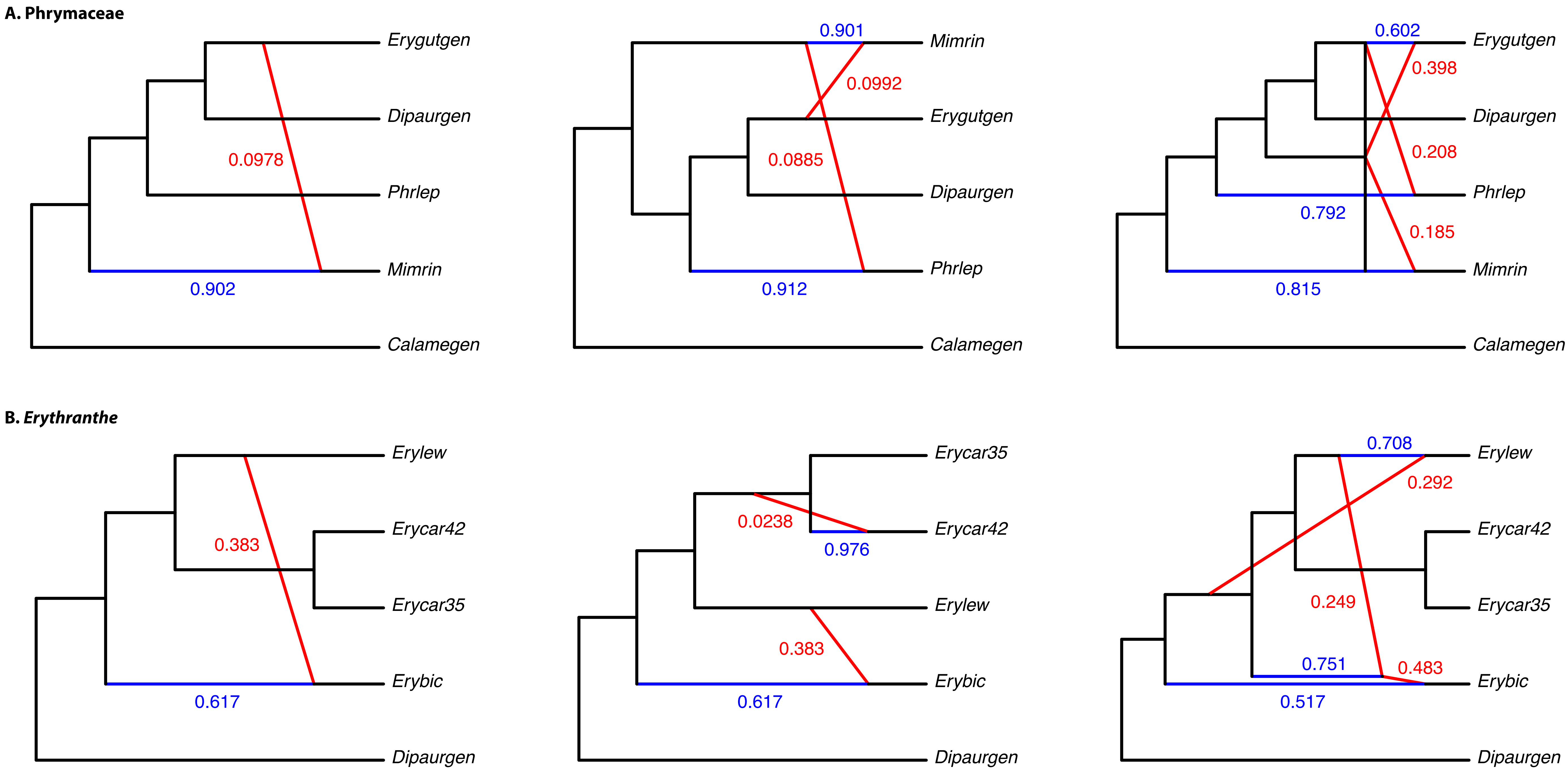

### Appendix S9

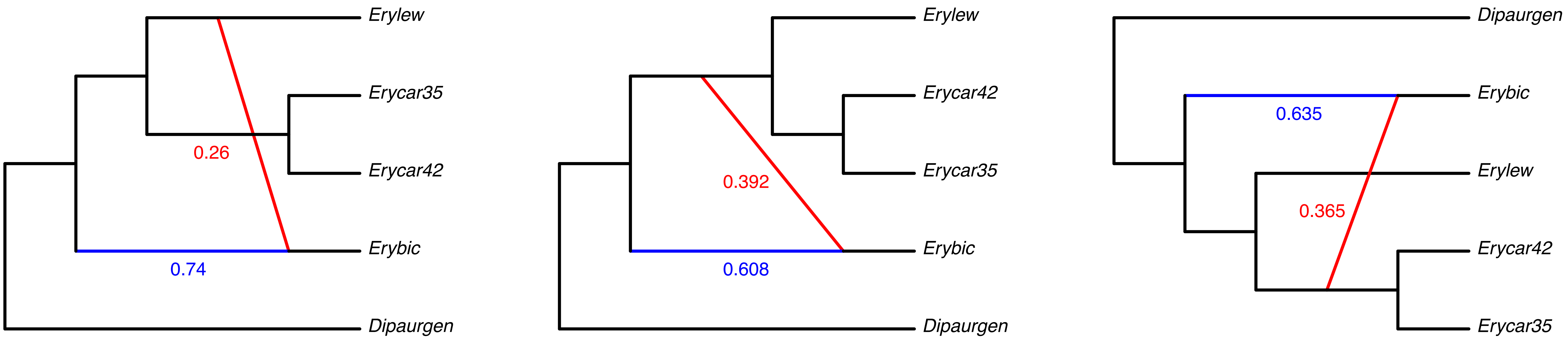

### Appendix S10

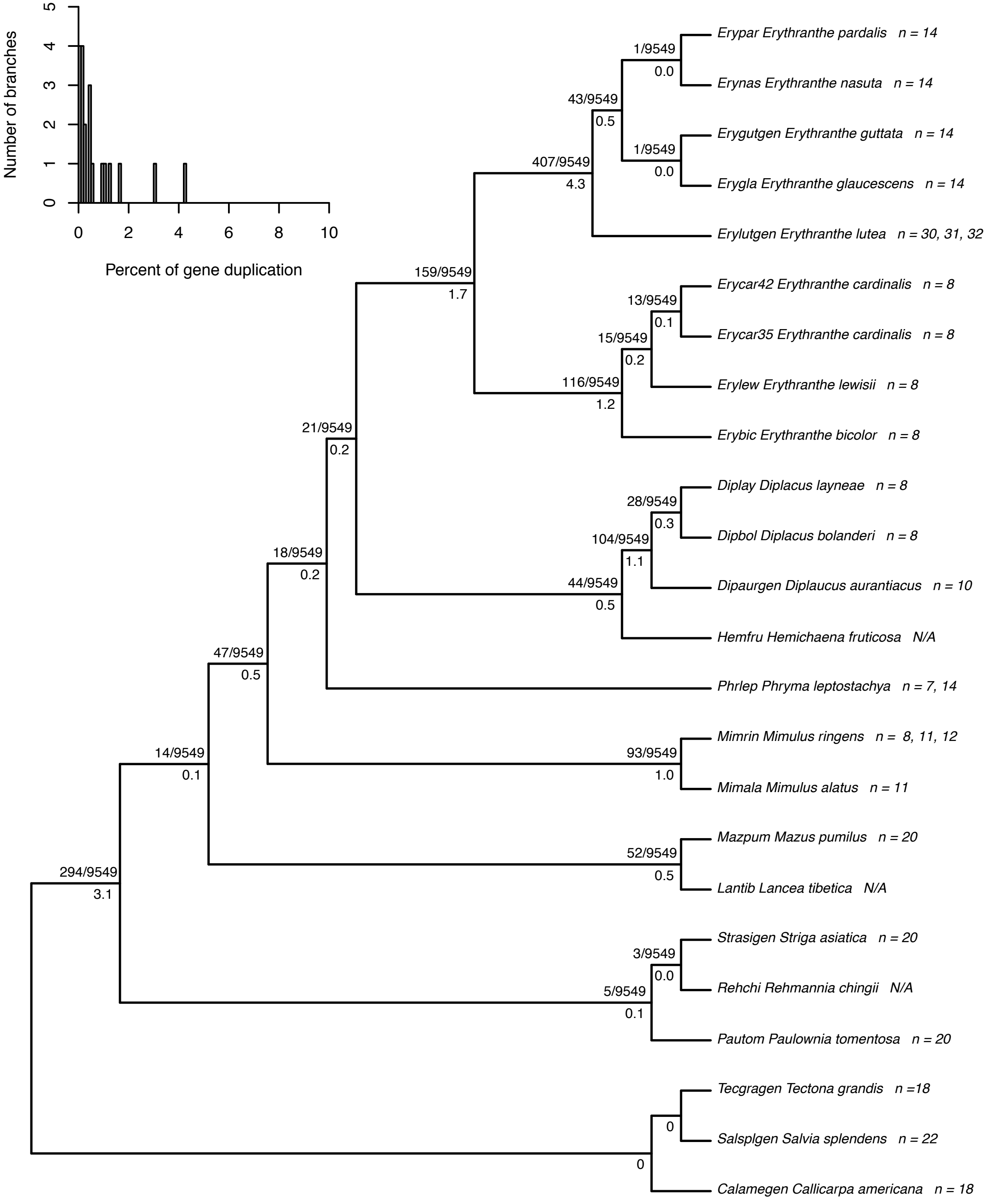

### Appendix S11

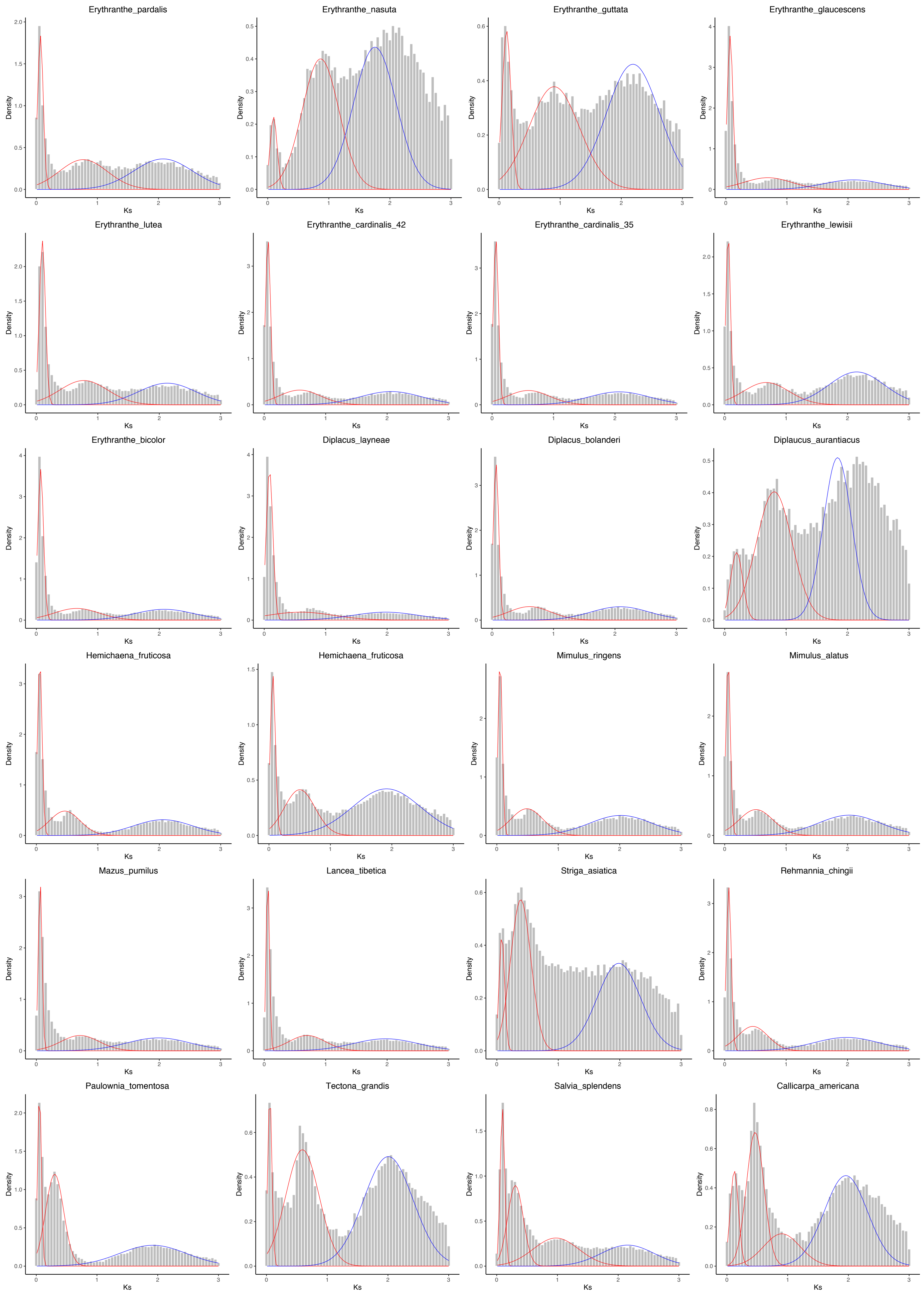

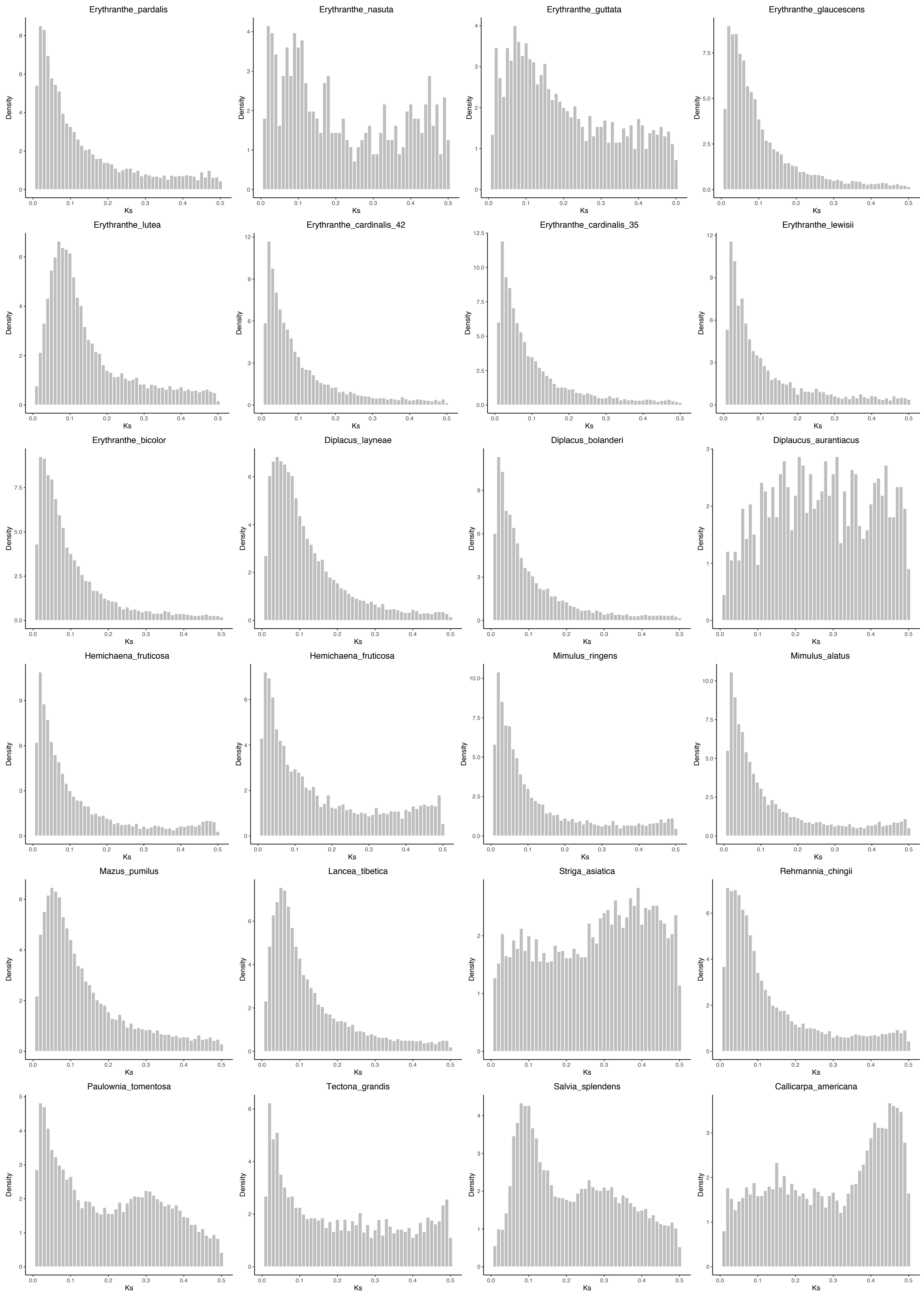

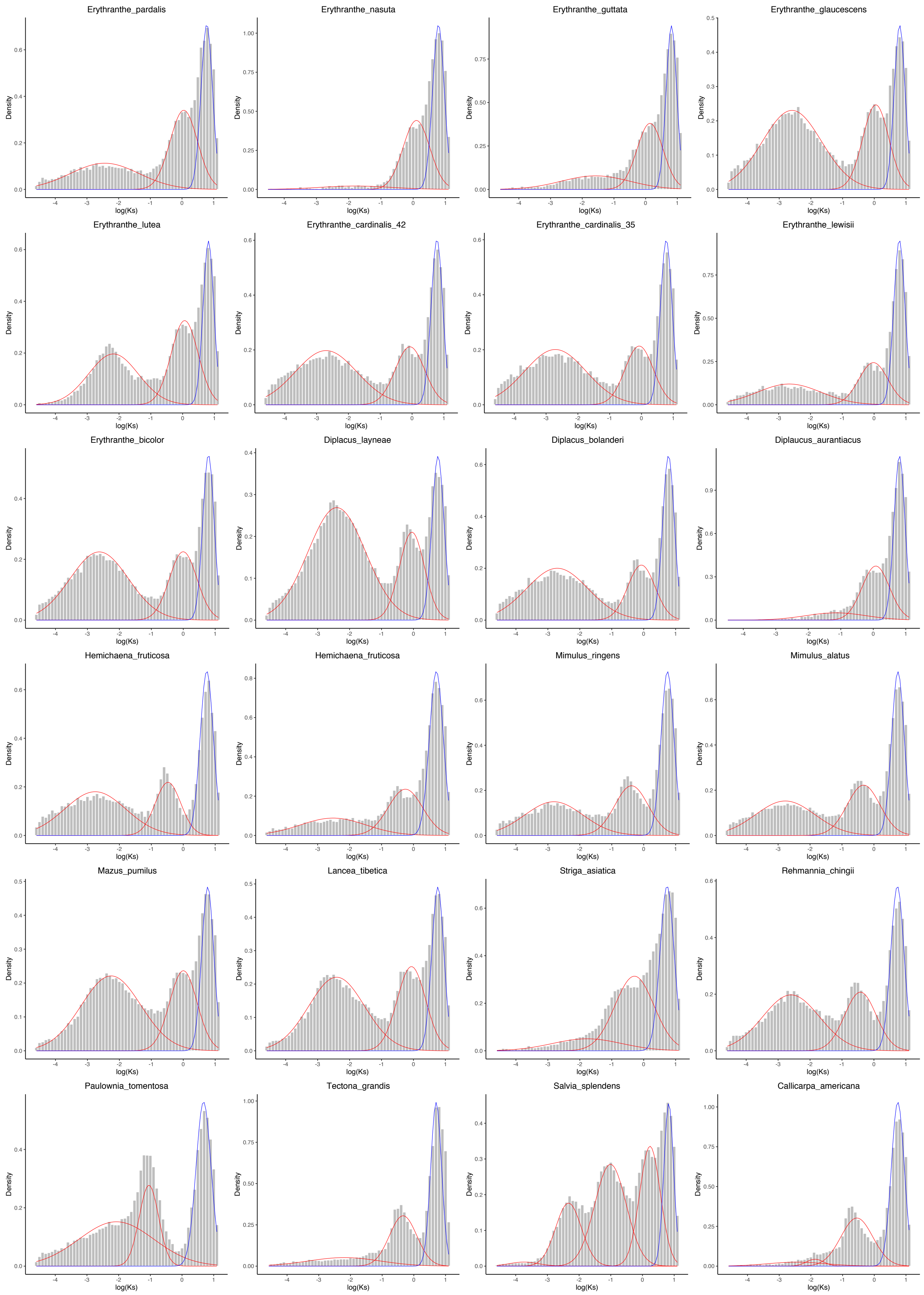
